# Attaching artificial Achilles and tibialis cranialis tendons to bone using suture anchors in a rabbit model: assessment of outcomes

**DOI:** 10.1101/2024.04.29.591695

**Authors:** Obinna P. Fidelis, Caleb Stubbs, Katrina L. Easton, Caroline Billings, Alisha P. Pedersen, David E. Anderson, Dustin L. Crouch

## Abstract

**Objective:** The purpose of this study was to investigate the factors associated with outcomes of attaching artificial tendons to bone using suture anchors for replacement of biological tendons in rabbits.

**Study Design:** Metal suture anchors with braided composite sutures of varying sizes (USP #1, #2, or #5) were used to secure artificial tendons replacing both the Achilles and tibialis cranialis tendons in 12 New Zealand White rabbits. Artificial tendons were implanted either at the time of (immediate replacement, n=8), or four weeks after (delayed replacement, n=4) resection of the biological tendon. Hindlimb radiographs of the rabbits were obtained immediately after surgery and approximately every other week until the study endpoint (16 weeks post-surgery).

**Results:** All suture anchors used for the tibialis cranialis artificial tendons remained secure and did not fail during the study. The suture linkage between the Achilles artificial tendon and anchor failed in 9 of 12 rabbits. In all cases, the mode of failure was suture breakage distant from the knot. Based on radiographic analysis, the mean estimated failure timepoint was 5.3±2.3 weeks post-surgery, with a range of 2-10 weeks. Analysis of variance (ANOVA) tests revealed no significant effect of tendon implantation timing or suture size on either the timing or frequency of suture anchor failure.

**Conclusion:** Based on the mode of failure, suture mechanical properties, and suture anchor design, we suspect that the cause of failure was wear of the suture against the edges of the eyelet in the suture anchor post, which reduced the suture strength below in vivo loads. Suture anchor designs differed for the tibialis cranialis and did not fail during the period of study. Future studies are needed to optimize suture anchor mechanical performance under different loading conditions and suture anchor design features.

## INTRODUCTION

A suture anchor consists of a suture and an anchor with an eyelet on the anchor through which the suture passes. Suture anchors are frequently used in human and veterinary musculoskeletal reconstruction to reconnect soft tissues, such as tendons and ligaments, to bones (Cho et al., 2021; Visscher et al., 2019). Suture anchors can offer quick, reliable, and stable fixation (Burnham et al., 2020; Ravin et al., 2005), and can achieve close apposition between the soft tissue and bone (Altiparmak & Uckan, 2013). An intimate interface between the anchor and suture is needed to ensure that the soft tissue is securely connected to the bone to facilitate healing (McFarland et al., 2005). Compared to other techniques for fixation of soft tissues to bone, the strength of suture anchors has been shown to be superior to other means of fixation such as bone tunnels, staples, screws, washers, or tapered plugs, (Barber et al., 1993).

Anchors can be classified as either screw-type or non-screw-type (Barber et al., 1995), with screw-type anchors having greater load-to-failure resistance compared with non-screw-type designs (Barber et al., 2006; Barber et al., 2003). The eyelets in screw-type anchors can be either raised above the screw or embedded within it. Anchors may also be bioabsorbable, nonmetallic, or metallic (Barber et al., 1993). Metallic anchors are relatively affordable with minimal undesirable biological reactions, ensuring safe and long-term fixation of the anchor within the bone while the tissue heals (Longo et al., 2019).

Mechanical failure of a suture anchor is a potential complication for clinical concern as it compromises the stability and integrity of the surgical reconstruction (Meyer et al., 2002). When failure occurs, the connection between soft tissue and bone becomes weak or ineffective, resulting in the need for revisions. Revision surgeries are often complex, costly, and burdensome for the patient. Sometimes, revisions may lead to more loss of bone tissue, depending on the nature of the reconstruction, as the bone is prepared to hold the anchor (Panegrossi et al., 2014). This highlights the importance of developing reliable and durable suture anchors to minimize the risk of failure. Several factors may affect the durability of suture anchors, including suture anchor design, material composition, and surgical techniques (Fleischli, 2018; Schanda et al., 2022).

Sutures associated with suture anchors tend to fail in one of two ways: at the knot, which is considered the weakest point of a tied suture, or at the eyelet of the anchor (Meyer et al., 2002). Using a model of bovine infraspinatus tendon repair, suture breakage at the eyelet was the most common mode of failure for metallic suture anchors with eyelets raised above the screw, when the anchor head was flush with the surface of the bone, and when the anchor head was raised above the bone (Bynum et al., 2005). Suture breakage also was the predominant mode of failure for two metallic screw type suture anchors evaluated in a human cadaveric shoulder using an arthroscopic Bankart method; although the study did not state if the sutures failed at the knot or around the eyelet (Barber et al., 1993). In an extensive study of the mechanical properties of suture-anchor interactions, fourteen different sutures were used in combination with thirty-one anchors; suture breakage at the midpoint of the tested strands was the failure mode for all the suture anchors (Barber et al., 2006; Barber et al., 2003).

In artificial tendon implant studies in New Zealand White (NZW) rabbits, we used screw-type metallic anchors, with the eyelet raised above the screw and placed at the point of insertion of the native tendons, to attach polyester suture-based artificial tendons used to replace the biological Achilles and tibialis cranialis tendons. In most of the study animals, the Achilles suture anchor failed prematurely, based on qualitative examinations of post-surgical radiographs; failures were confirmed by ex vivo dissection. The failures were unexpected because the reported knot strength of the largest sutures used (#5 FiberWire) was 295N (Arthrex Inc, 2014), which far exceeded the reported peak in vivo Achilles tendon force of 57.7 N during hopping gait in NZW rabbits (West et al., 2004). Toward determining the cause of the suture anchor failures in the rabbits, the objective of this study was to estimate and determine the timing and mode of failure, respectively. The results of our study will inform the design and selection of suture anchors for clinical applications.

## MATERIALS AND METHODS

### Artificial tendon fabrication

We fabricated custom polyester suture-based artificial tendons to replace the biological Achilles and tibialis cranialis tendons, as previously described (Hall et al., 2023). The artificial tendon design was adapted from an existing device (Melvin et al., 2010; Melvin et al., 2012) using customized USP size 0 braided polyester sutures (RK Manufacturing Corp, Danbury, CT, USA). The sutures were cut to a length of 12 inches, double-armed with swaged 3/8-circle tapered point needles (0.028-inch wire diameter). Artificial Achilles tendons were made using three suture strands that were folded in half to form a loop at the mid-point, then braided to the desired length, yielding 6 suture threads for muscle attachment; tibialis cranialis artificial tendons included 2 suture strands, folded in half to form a loop, then braided to the desired length yielding 4 suture threads for muscle attachment. To prevent tissue adhesion, medical-grade biocompatible silicone (BIO LSR M340, Elkem Silicones, Lyon, FR) was applied to the braided portion of the tendon. The tendons were then cleaned and sterilized in preparation for surgery.

### In Vivo Study

All animal procedures were approved by the Institutional Animal Care and Use Committee at the University of Tennessee, Knoxville. The study included twelve female New Zealand White rabbits with an average age of 19.9 ± 2.17 weeks and an average body mass of 3.52 ± 0.35 kg at the time of first surgery. All rabbits were part of a larger study to determine the effect of tendon replacement timing, immediate versus delayed (5 weeks), on hindlimb movement biomechanics. Surgeries were performed on multiple days over a 35-week period (Figure 1) under general anesthesia and the rabbits were given pain management with bandaging of the operated limb, post-surgery. First, the biological Achilles and tibialis cranialis tendons were excised from the musculotendinous junction to the tendon enthesis. Artificial Achilles and tibialis cranialis tendons were then implanted either at the time of (immediate) or 5 weeks after (delayed) biological tendon excision (Table 1). The proximal ends of the artificial tendons were sewn into the distal end of the respective muscle. The distal ends of the artificial tendons were attached near the respective biological tendon insertion points on the bone using metallic screw-type anchors with eyelets located within a raised post. The anchor (Figure 2) for the artificial Achilles tendon (part no. 60-27-09, 2.7 mm x 9 mm, IMEX Inc., Longview, TX) was inserted in a proximal to distal angle through the cranial aspect of the calcaneus close to the point of insertion of the biological tendon. In one of the rabbits (R4), the tendon of the superficial flexor digitorum excised and replaced instead of the Achilles tendon. Either USP size #1, #2 or #5 braided composite suture (Fiberwire, Arthrex Inc., Naples, FL) was passed through the eyelet for use to secure the artificial tendon to the anchor. The anchor for the tibialis cranialis artificial tendon (2 mm x 6mm, Jorgensen Laboratories LLC., Loveland, CO) was inserted in the proximal, dorsal aspect of the 2^nd^ metatarsus (MT II) and accompanied with either a size #1 or #2 suture.

**Table 1.**
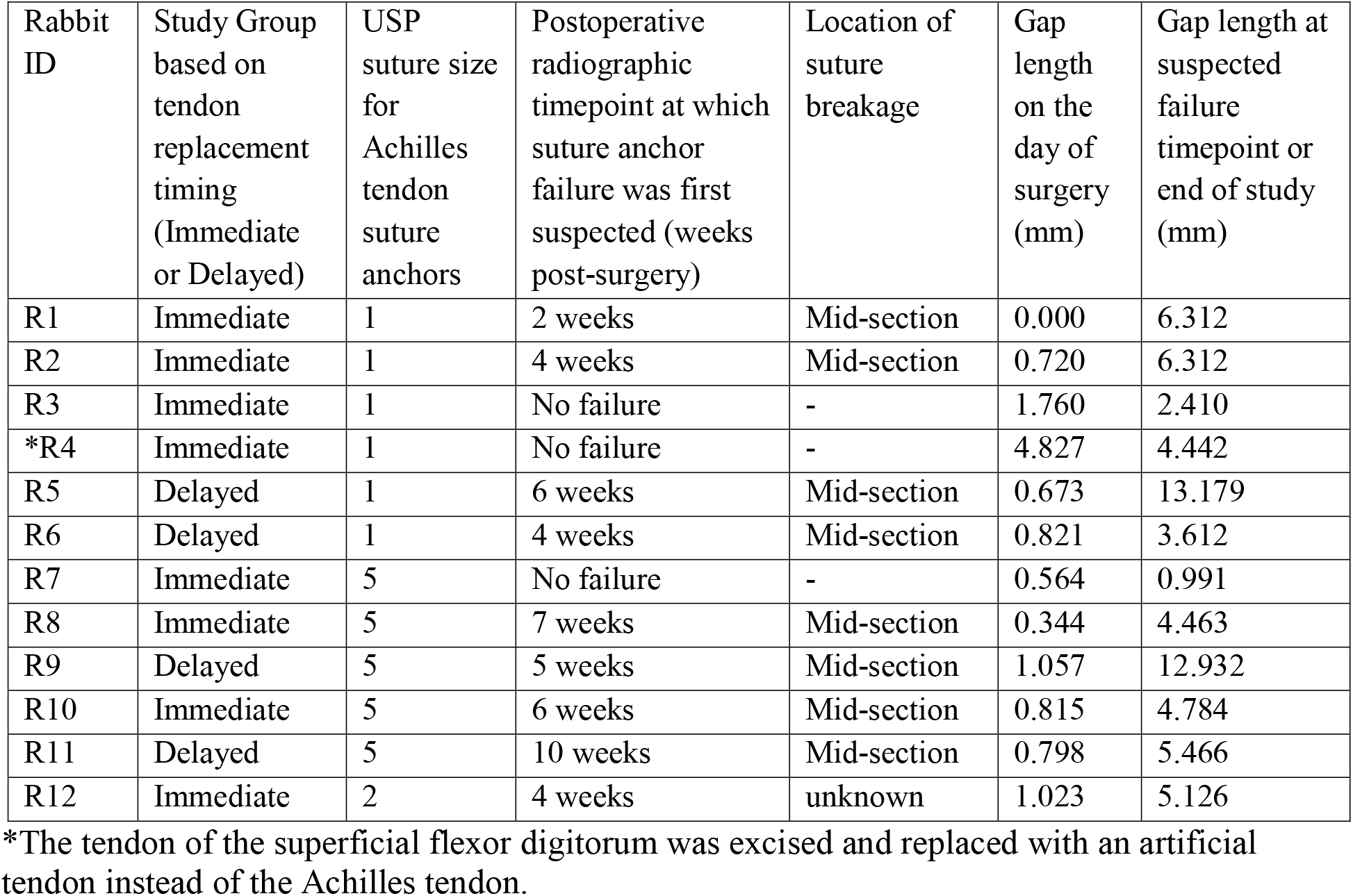
Summary of results for Achilles artificial tendon suture anchors.

**Figure 1.**
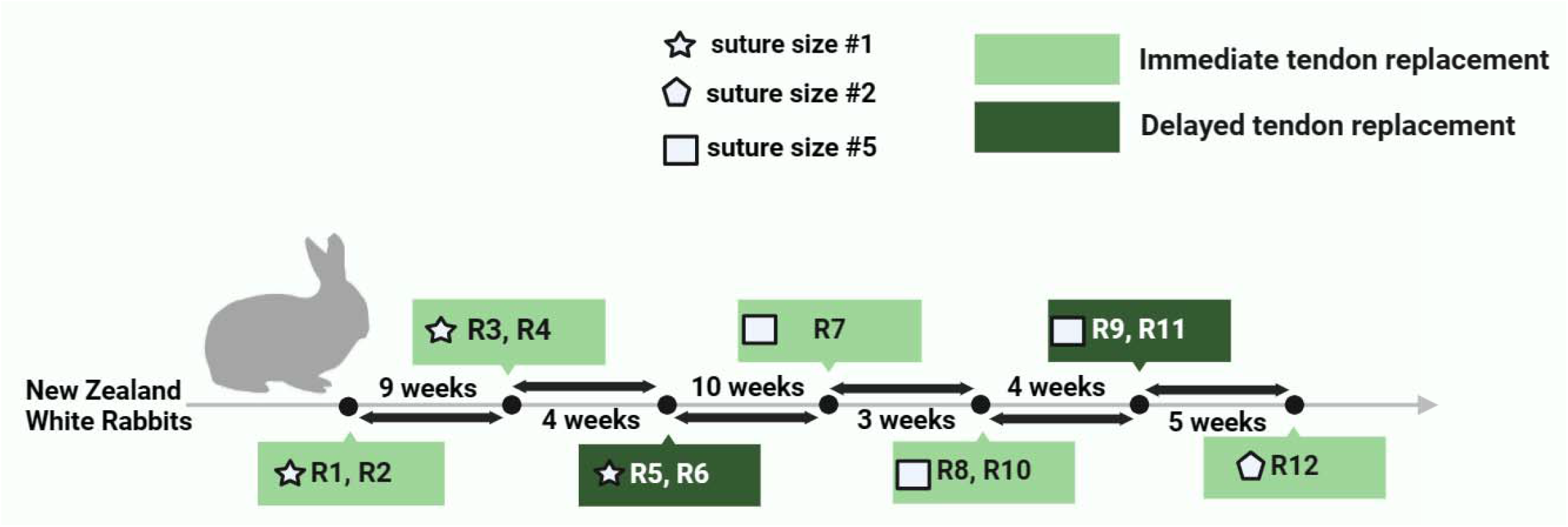
Timeline of surgeries in the study showing the tendon replacement timing (immediate or delayed) and the suture size used for the Achilles tendon suture anchor for rabbits R1 through R12.

**Figure 2.**
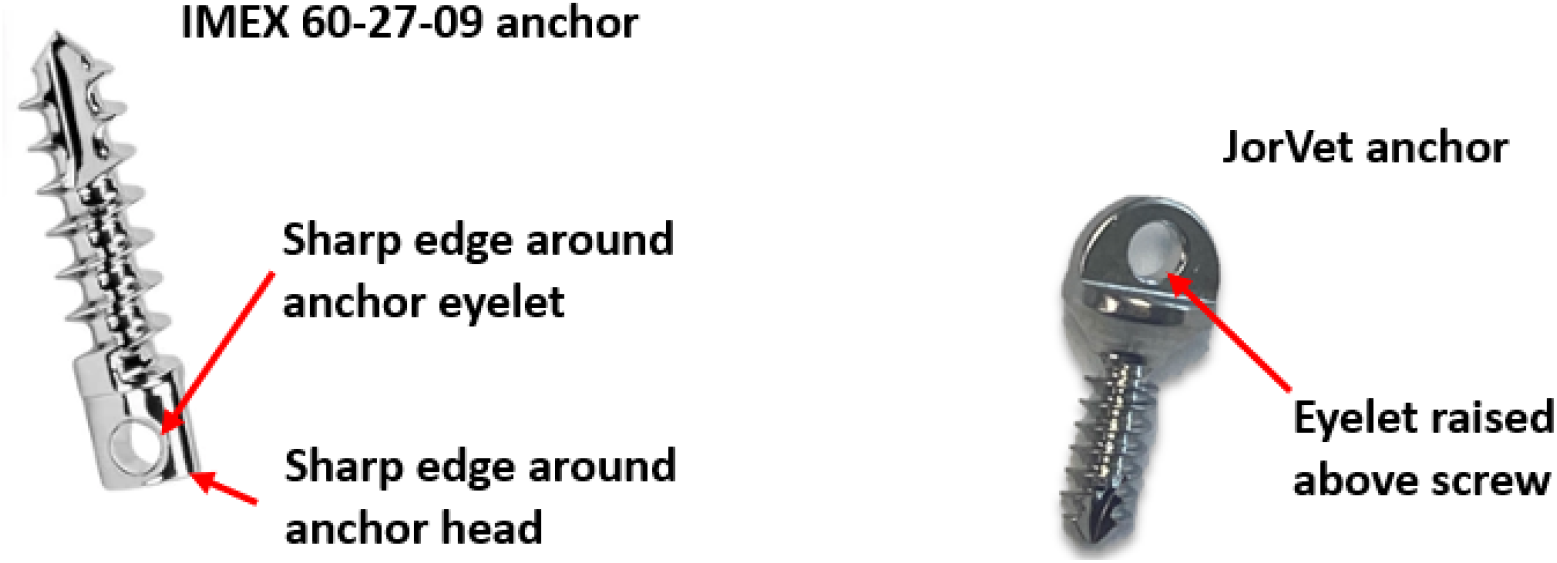
IMEX 60-27-09 anchor (left) used to attach artificial Achilles tendon. A second raised eyelet design (JorVet 2 mm x 6 mm) used to attach artificial tibialis cranialis tendon is also shown (right).

Radiographs of the operated limbs were taken at the time of surgery and approximately once every other week post-surgery to monitor the placement and integrity of the suture anchors and artificial tendons. Both lateral and cranial-caudal views were done at each timepoint.

At the end of the study (16 weeks post-surgery), the rabbits were euthanized. The hindlimbs were collected via disarticulation of the limb at the level of the hip joint, fixed in 10% phosphate-buffered formalin for at least 5 days, and then stored in 70% ethanol. Following fixation, the hindlimbs were dissected to permit visual inspection of the suture anchor and confirmation of suture breakage in all rabbits. We also determined the mode of suture failure, either at the knot or at the mid-section (i.e., away from the knot).

### Data Analysis

The artificial tendon and metallic anchor were visible on radiographs, but not the connecting suture. Thus, failure of the tendon suture anchors by suture breakage was suspected when, qualitatively, the gap length between the artificial tendon and anchor was substantially longer than that observed immediately after surgery. From the digital radiographs, we used image processing software (ImageJ, NIH) to measure the tendon-anchor gap length, defined as the length from the head of anchor to the most distal visible point on the artificial tendon. No tibialis cranialis suture anchor failures occurred; thus, gap lengths were only measured for the Achilles tendon suture anchors. Post-surgery gap lengths were compared to those at the time of surgery to estimate the timing of suture anchor failure, defined as the first radiographic timepoint at which gap length had increased substantially (> 5 mm) compared to the previous timepoint with no return to the previous shorter gap length.

The estimated time of Achilles tendon suture anchor failure was used to perform a survival analysis to determine the probability of failure as a function of time post-surgery. For all samples, we defined suture-anchor failure as the timepoint at which a large gap distance was detected radiographically. The exact time of failure could not be determined because of the pre-determined radiography timepoints. The number of events, cumulative survival, and cumulative failure (1 – cumulative survival) was calculated for each time point using equation 1.

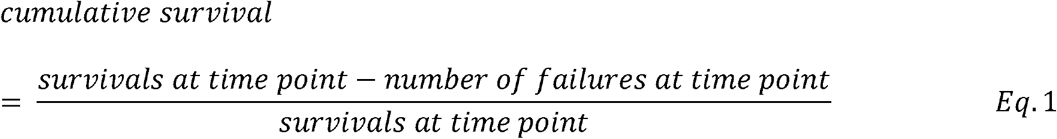

The survival distribution function was calculated by multiplying the cumulative survival rates and plotted against time to obtain a Kaplan-Meier survival curve. We applied the fundamental assumptions of the Kaplan-Meier survival analysis (Etikan et al., 2017).

A two-way analysis of variance (ANOVA) was used to determine the potential effect of two key variables, surgery type (immediate and delayed tendon replacement) and suture size (#1, #2, and #5), on the timing of suture anchor failure. Additionally, the possible effect of surgery type and suture size on the *frequency* of suture anchor failure was investigated using a Fisher’s exact test with the data grouped as *number of failures* associated with each surgery type and with each suture size. For all statistical comparisons, a p-value of < 0.05 was considered significant.

## RESULTS

Radiographs were inspected qualitatively to identify suspected failures of suture anchors. Based on this inspection, in vivo failure of the suture anchor used to attach the artificial Achilles tendons was initially suspected in 9 of the 12 rabbits (Figures 3 and 4). Conversely, there was no qualitative evidence of failure of the suture anchors used for attaching the tibialis cranialis artificial tendons.

**Figure 3.**
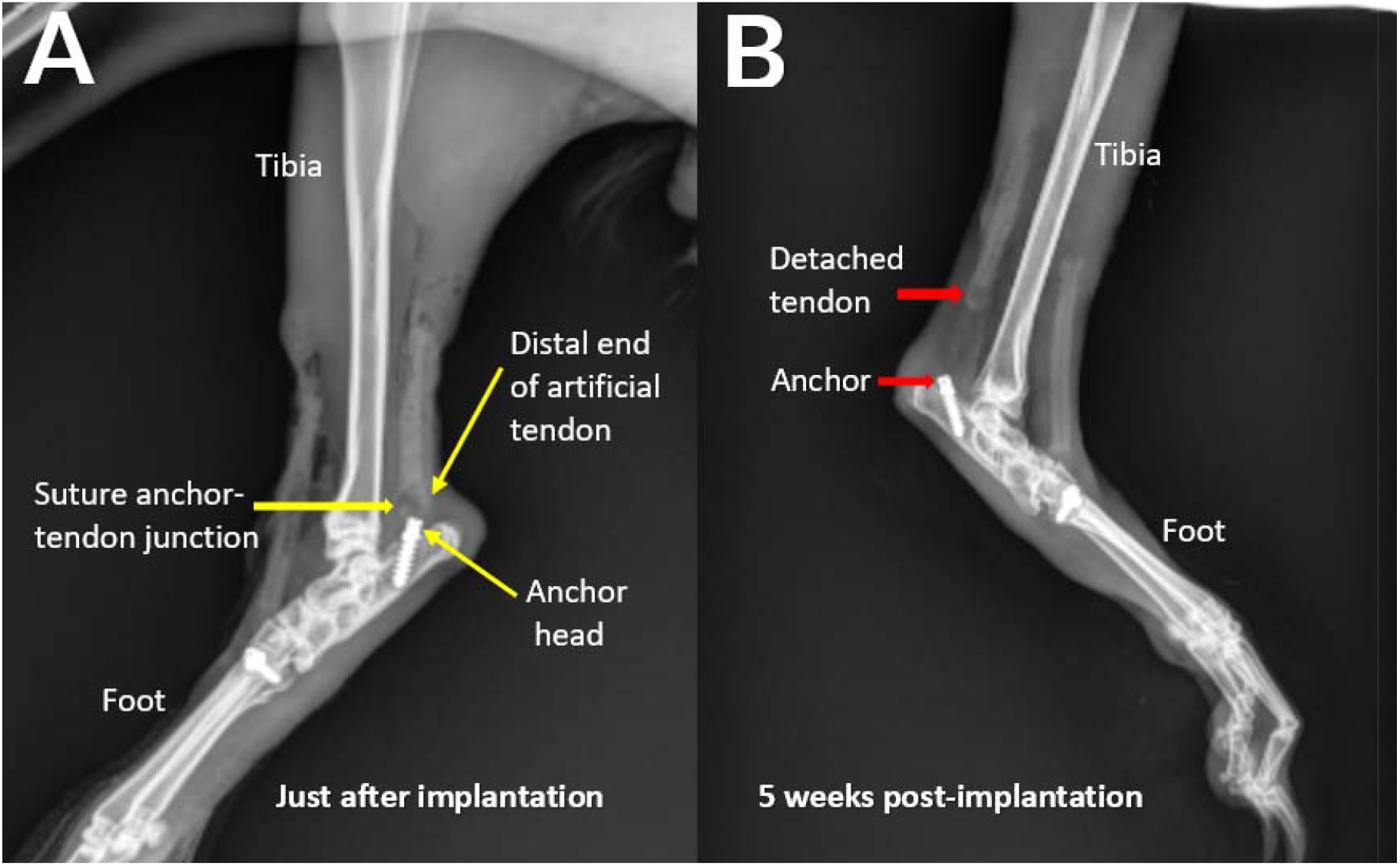
Exemplary hindlimb radiographs showing attachment of the artificial tendon just after surgery and detachment of the tendon from the anchor following failure of the suture anchor after five weeks for rabbit R9.

**Figure 4.**
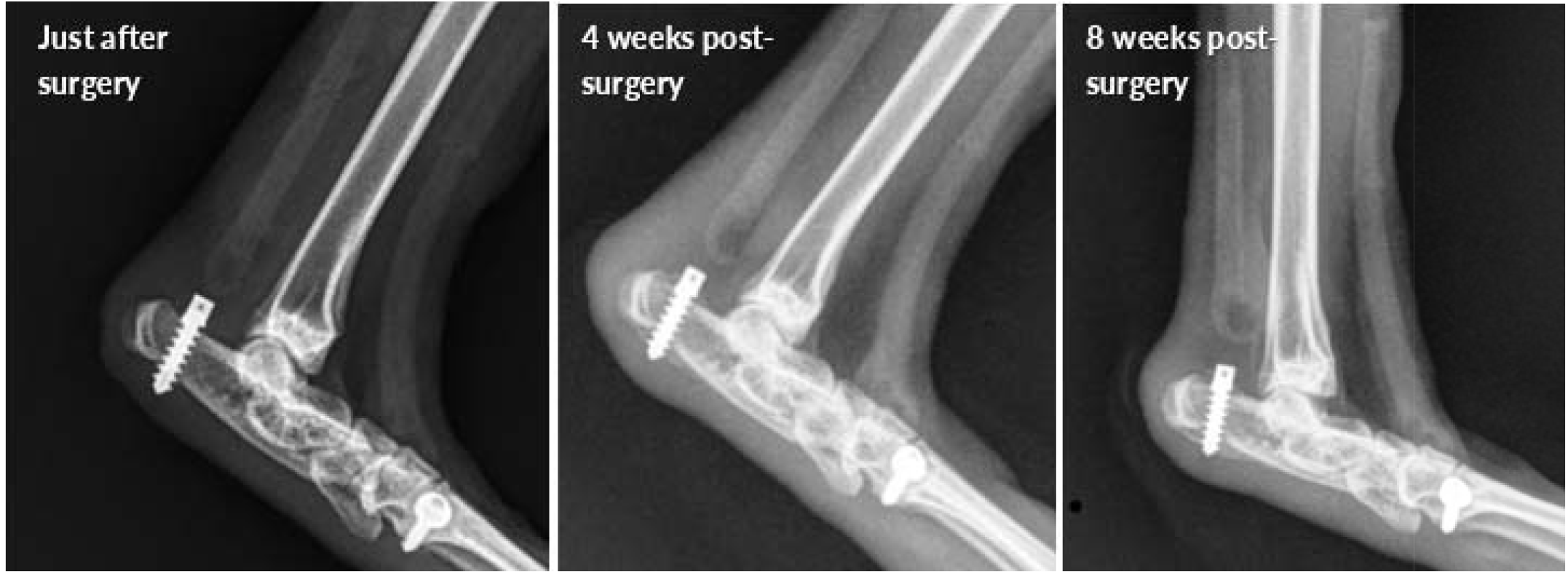
Hindlimb radiographs from rabbit R11 showing a gradually increasing gap length between the anchor and Achilles artificial tendon.

All specimens were inspected post-mortem by gross dissection. For all available specimens, the failure states (failed or not) of all suture anchors determined by post-mortem dissection were consistent with those determined radiographically (Figure 5). For all failure cases, the mode of suture anchor failure was suture breakage at the mid-section, i.e., away from the knot (Table 1).

**Figure 5.**
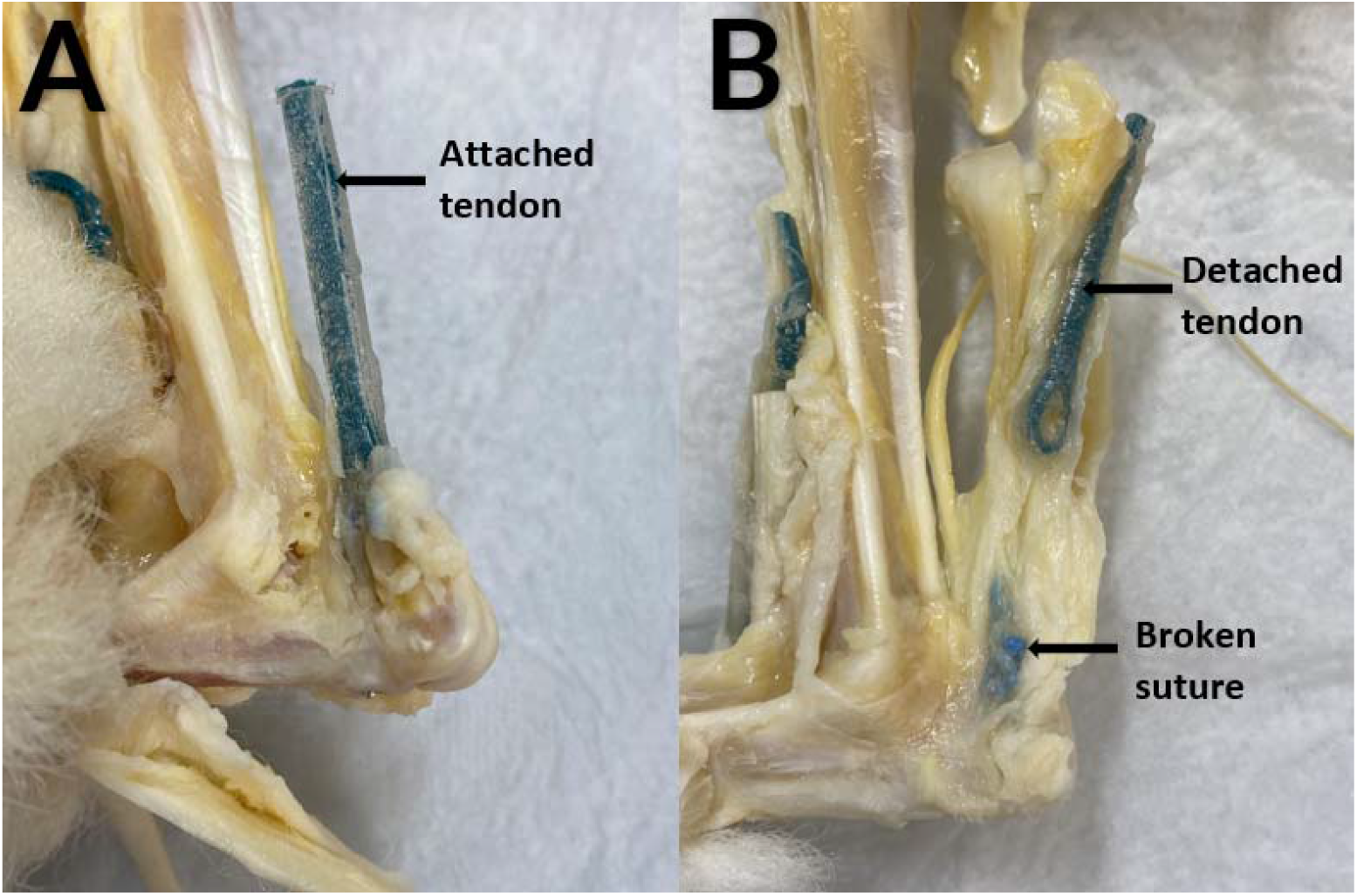
(A) Artificial Achilles tendon securely attached to the calcaneus by an intact suture anchor (B) Artificial Achilles tendon detached from a broken suture in a rabbit.

For the nine Achilles tendon suture anchor failures, we measured a substantial (≥5mm) increase in gap length between two adjacent timepoints, with no return to the pre-failure length (Figure 6, Table 1). The timepoint at which the substantial increase in gap length was first observed was designated as the failure timepoint. The earliest failure (R1) occurred only 2 weeks after implantation. Three suture anchors failed at 4 weeks post-implantation (R2, R6, and R12), one at 5 weeks (R9), two at 6 weeks (R5 and R10), and one each at 7 weeks (R8) and 10 weeks (R11). In three rabbits (R3, R4 and R7), the suture anchor did not fail. However, it should be noted that the tendon of the superficial flexor digitorum was replaced in R4, instead of the Achilles tendon. In some rabbits, gap length continued to increase over time after the suspected failure timepoint. Of the nine suture anchors that failed, the mean failure timepoint was 5.3±2.3 weeks post-surgery, with a range of 2-10 weeks.

**Figure 6.**
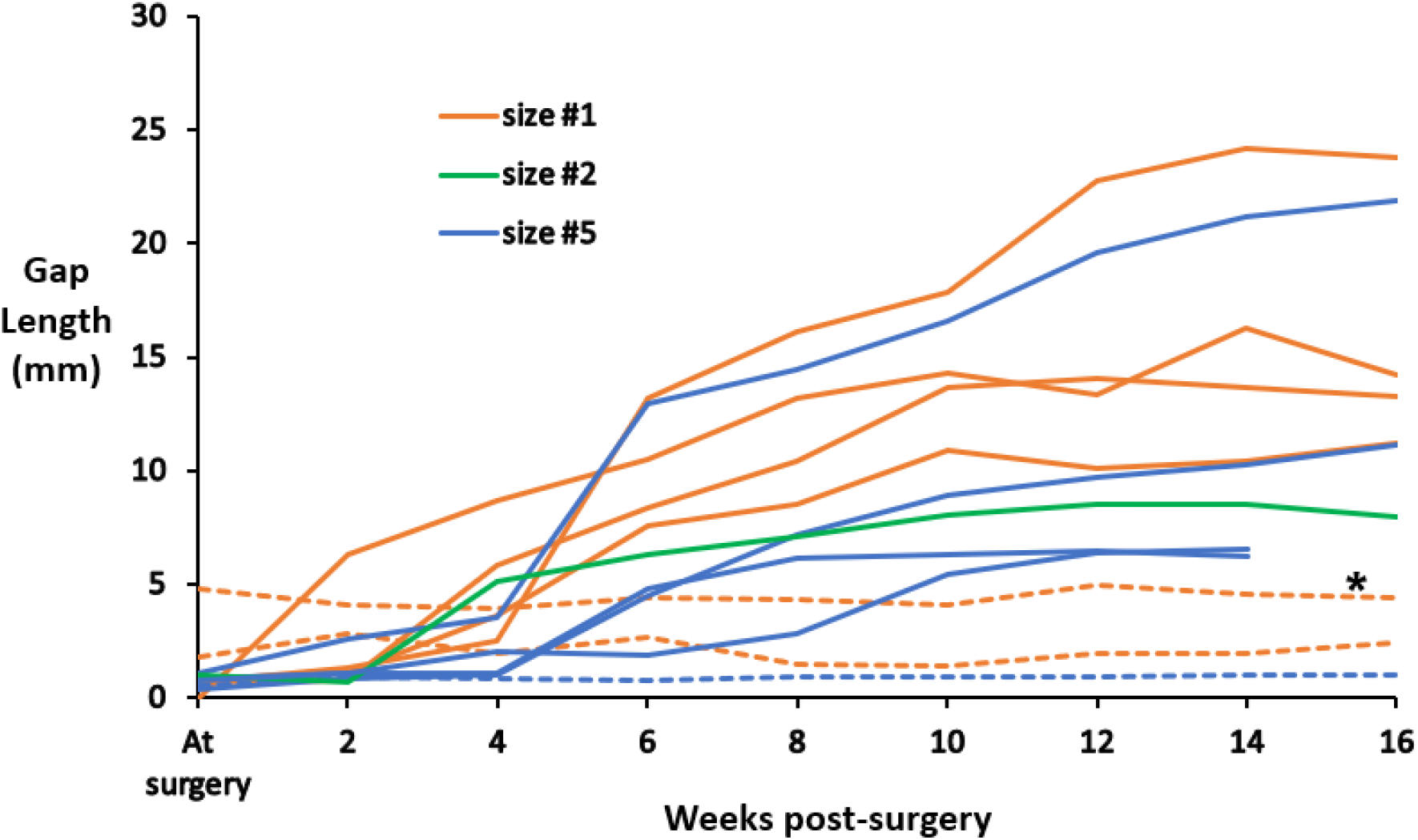
Gap length between the anchor and Achilles artificial tendon as measured from radiographs. In nine rabbits, the gap length increased by at least 5mm between two consecutive radiographic timepoints (solid lines); these rabbits were suspected to have suture anchor failure. The gap length remained about the same in three rabbits (dashed lines); these were suspected to have intact suture anchors throughout the entire study. *In R4, the tendon of the superficial flexor digitorum was replaced, instead of the Achilles tendon.

The Kaplan-Meier survival analysis (Table 2, Figure 7) revealed that 92 percent of the suture anchors survived beyond the initial two weeks. As time progressed, the survival rate decreased gradually, with 67 percent of the anchors surviving beyond 4 weeks, 58 percent of the anchors surviving beyond 5 weeks, 42 percent surviving for 6 weeks, and 33 percent of the suture anchors surviving beyond seven weeks. Finally, 25 percent of the anchors survived up to the end of the study at 16 weeks.

**Table 2:** Kaplan-Meier survival rate and distribution function for 12 suture anchors.

| Time when failure occurred (weeks) | Number of failed suture anchors | Number of censored data | Survival rate of suture anchors | Survival distribution function |
| --- | --- | --- | --- | --- |
| 0 | 0 | 0 | 1.0000 | 1.0000 |
| 2 | 1 | 0 | 0.9167 | 0.9167 |
| 4 | 3 | 0 | 0.7272 | 0.6667 |
| 5 | 1 | 0 | 0.8750 | 0.5833 |
| 6 | 2 | 0 | 0.7140 | 0.4165 |
| 7 | 1 | 0 | 0.8000 | 0.3332 |
| 10 | 1 | 0 | 0.7500 | 0.2499 |
| 16 | 0 | 3 | 1.0000 | 0.2499 |

**Figure 7.**
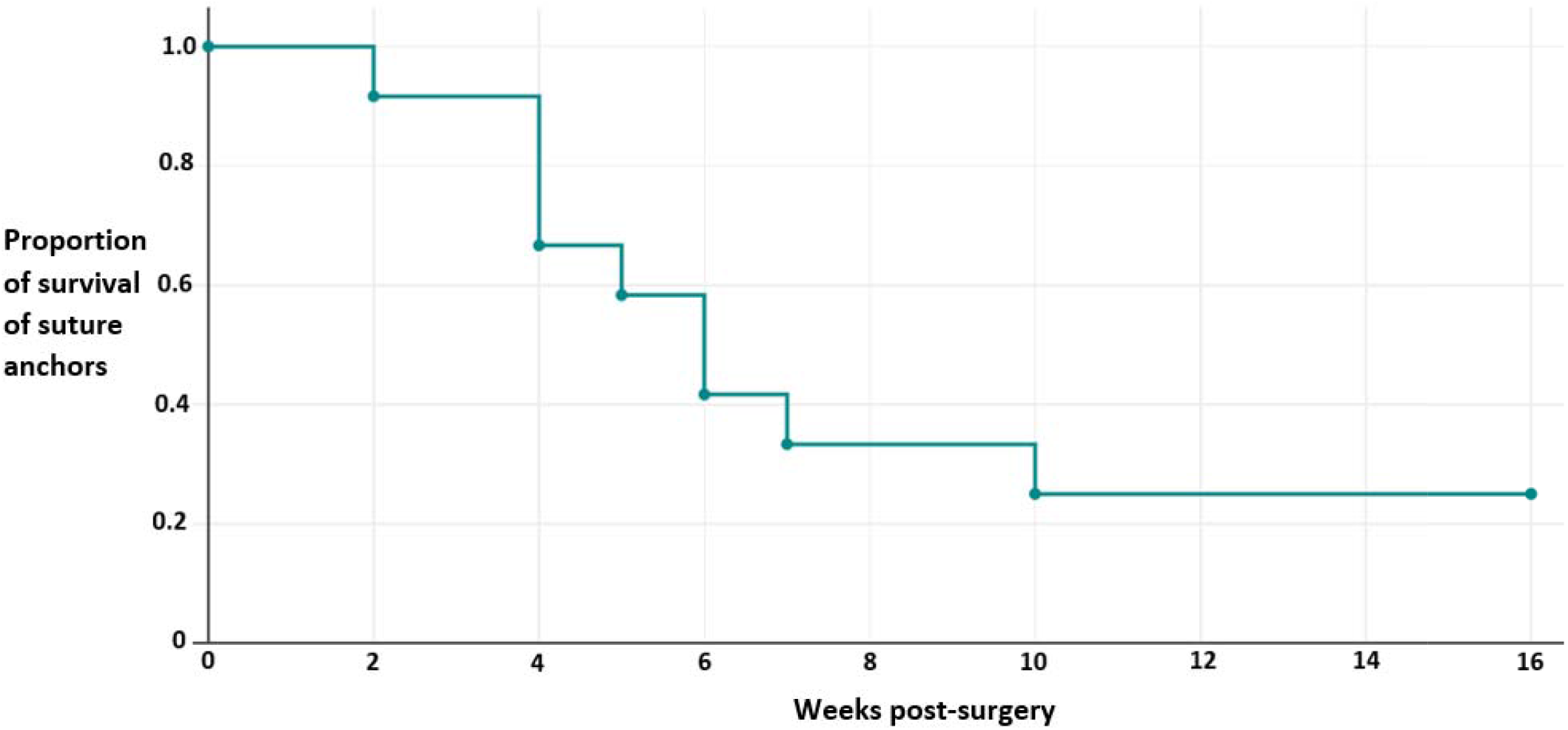
Kaplan-Meier survival curve showing the probability of suture anchor failure at the respective time intervals.

Analysis of variance (ANOVA) showed that the timing of failure was not significantly affected by either the timing of surgery or the size of the suture. Similarly, Fisher’s exact test indicated that there were no statistically significant differences in the frequency of failure across the different surgery types or suture sizes.

## DISCUSSION

In our rabbit model, bone suture anchors were successful in securing an artificial tendon implant to the point of insertion of the native tendon. The frequent failures of the Achilles suture anchor-tendon implant linkage were surprising and characterized by a sudden (within two-week period between consecutive radiographic timepoints) and large increase in gap length, followed by gradual continued increase in gap length. The gradual increase in gap length may have been due to shortening of the muscle fibers (Maffulli, 1999; Meyer, Farshad, et al., 2011). Muscles retract after tendon ruptures, with the degree of retraction depending on the muscles involved and the severity of the rupture(Gil-Melgosa et al., 2021; Meyer, Gerber, et al., 2011). Mean gap length of up to 6.0 cm has been reported in cases of distal biceps tendon ruptures (Samra et al., 2019) and up to 4.8 cm gap length in cases of supraspinatus tendons ruptures (Meyer, Farshad, et al., 2011). The implications of muscle retraction include muscle atrophy, loss of muscle mass and strength, muscle weakness and a loss of tissue quality and quantity, which increase the risk of complications and decrease the success rate of surgical repair (Lakemeier et al., 2010).

The location of suture failure provides important clues about the cause of failure. The knot is considered the most severe stress point with sutures, and therefore, a weak link in a suture anchor. Therefore, failures at the knot are usually attributed to excessive loading. Conversely suture anchor failure away from the knot suggests wear or abrasion of the suture against part of the anchor. This mode of failure can reduce suture strength by as much as 73% (Meyer et al., 2002). The reduction in strength was presumably more pronounced in our rabbit model when using size #5 sutures, given that the reported in vivo Achilles tendon loading (57.7 N) was 80% lower than the reported strength of the suture (295 N).

Suture anchor design, especially the design of the eyelet through which the suture passes, likely played a significant role in the observed failures. The eyelets of suture anchors are generally configured in two ways: either raised above the screw or embedded within the screw. Anchors with raised eyelet designs offer greater versatility because they can accommodate sutures of various sizes. However, this design has drawbacks; a raised eyelet allows relatively unrestricted freedom of suture movement through the eyelet. Conversely, the embedded eyelet design (e.g., Mini Corkscrew AR-1319FT, 2.7 mm x 7 mm, Arthrex Inc., Naples, FL, and the Anika 2.5 mm MiTi, 2.5 mm X 5.7 mm, Parcus Medical, LLC, FL), while limiting the range of suture sizes that can be accommodated, restricts suture motion through the eyelet. Therefore, sutures may wear more rapidly in suture anchors with raised eyelets than with embedded eyelets; future studies should quantify the effect of suture motion on the extent and rate of suture wear.

With either raised or embedded eyelets, another factor that likely affects suture failure is the sharpness of the eyelet edges. Here we define sharpness as the radius of a corner feature, where a lower radius corresponds to a sharper edge; previous finite element simulations have shown a strong relationship between cutting force and edge radius, though in a different context (blade cutting into soft material) (McCarthy et al., 2010; Schuldt et al., 2016). One study noted that, due to the small size of suture anchors, many have eyelets with sharp edges (Meyer et al., 2002), which may cause the suture to wear more rapidly. The suture anchor used in the rabbits had two edges that could be both considered sharp and contacted by the suture in vivo: the edges around the eyelet and the edge around the oval-shaped top of the screw (Figure 2). In previous mechanical tests of suture anchors, edge radius or other potential aspects of sharpness (e.g., wedge angle (McCarthy et al., 2010) were not measured. Future studies should quantify the effect of edge sharpness on suture failure conditions, such as failure load, to inform the design of suture anchors.

Despite having a similar eyelet configuration as the Achilles tendons suture anchor but smaller suture, the tibialis cranialis suture anchors did not fail in any of the 12 rabbits. There are several factors that potentially explain the difference in failure rates between the suture anchors for the two different artificial tendons. For one, the Achilles tendon is expected to experience greater biomechanical loads than the tibialis cranialis tendon. The triceps surae-Achilles tendon unit provides body weight support against gravity and propels the body during locomotion (Machado et al., 2021); conversely, the tibialis cranialis muscle-tendon unit contributes to foot motion during the swing phase of locomotion (Viidik, 1969). In vivo motion of the artificial tendon and suture relative to the anchor, which could affect wear of the suture against the anchor eyelet, could have been greater for the Achilles tendon than for the tibialis cranialis tendon. Finally, the eyelet design and material characteristics, including the radii (sharpness) and smoothness of edges, could have been more averse to suture durability for the Achilles tendon anchor, but this could be verified in future studies using optical profilometry.

The results of the ANOVA and Fisher’s exact tests provided further evidence to suggest that the suture breakage was caused in part by wear or abrasion of the suture against the anchor rather than exclusively due to biomechanical overload. These statistical tests found no effect of suture size on the timing or rate of failure. If overloading was the cause of failure, one may expect that failures would occur possibly later or at a lower rate for anchors with larger suture. This is especially true given that, as previously mentioned, the expected in vivo load (60 N) was substantially less than the strength of the largest suture used (295 N). Additionally, the ANOVA and Fisher’s tests indicated no effect of the type of surgery (immediate vs. delayed replacement); this result suggests that the failure conditions were similar between the two surgery types.

In the present study, the survival analysis provided information about the likelihood of failure for the suture anchor used in the study along with their corresponding survival rates. The survival analysis of the suture anchors showed that only about 25 per cent of the suture anchors survived up to 16 weeks post implantation, as verified by postmortem dissection of the limbs. These findings provide valuable insights into the durability and performance of the suture anchors over time, showing the proportion of anchors that effectively withstood the loading conditions in the rabbits post-implantation and for the duration of the study. By incorporating both tabular and graphical representations, this analysis offers a comprehensive understanding of the survival patterns and outcomes of the suture anchors under investigation. It will also assist in making important decisions about our choice of suture anchors for future research.

Suture breakage due to wear is partly a function of the number of loading cycles (Bardana et al., 2003). A limitation of our investigation is that the number of loading cycles that the suture anchors accumulated in vivo is unknown. Non-standard radiographic positioning also can play a role in the gap measurement. As noted in the method, one rabbit had the tendon of the superficial flexor digitorum replaced instead of the Achilles tendon. It is unclear if this would have affected the magnitude of loading of the artificial tendon because the artificial tendon did not fail in this rabbit. The range of timepoints over which failure occurred (2-10 weeks post-surgery) as well as the standard deviation of the mean failure timepoint (standard deviation was 43% of the mean) were both relatively large. Additionally, the earliest failure timepoint (2 weeks post-surgery) may seem too soon for wear to have occurred. However, previous in vitro mechanical tests observed that, for some suture anchors and loading conditions, the number of loading cycles to failure had similar variability and was low (<10 cycles) (Bardana et al., 2003). The number of loading cycles could be estimated in future in vivo studies by, for example, using wearable motion sensors to estimate the number of “steps” the animal takes.

## CONCLUSION

Suture anchors used to attach an artificial Achilles tendon to bone failed in 9 of 12 rabbits; no failures were observed for a different suture anchor used to attach tibialis cranialis artificial tendons. All failures of the Achilles tendon suture anchor were characterized by suture breakage at the mid-section (i.e., away from the knot). Based on this failure mode and other aforementioned factors, we suspect that the failures were caused by wear of the suture against an edge of the suture anchor, which reduced the suture strength below in vivo loads. The failures may be partly attributed to the anchor design and to loading conditions. Therefore, the next steps in our study include measurement of (1) suture anchor design features such as edge radii and (2) in vitro strength of different suture anchors under physiologic loading conditions.

## Acknowledgements

Thanks to the Office of Laboratory Animal Care staff for assisting with surgeries. Thanks to the animal housing facility staff for assisting with animal care. Thanks to Elizabeth Croy for setting up operating room and surgery supplies.

